# Identification of the aldolase responsible for the production of 22-hydroxy-23,24-bisnorchol-4-ene-3-one from natural sterols in *Mycolicibacterium smegmatis*

**DOI:** 10.1101/2023.02.10.527981

**Authors:** Gabriel Hernández-Fernández, Miguel G. Acedos, José L. García, Beatriz Galán

## Abstract

Mycobacterial mutants blocked in ring degradation constructed to achieve C19 synthons production, also accumulate by-products such as C22 intermediates throughout an alternative pathway reducing the production yields and complicating the downstream purification processing of final products. In this work, we have identified the *MSMEG_6561* gene, encoding the only aldolase on the chromosome responsible for the transformation of 22-hydroxy-3-oxo-cholest-4-ene-24-carboxyl-CoA (22-OH-BCN-CoA) into the 22-hydroxy-23,24-bisnorchol-4-ene-3-one (4-HBC) precursor (20S)-3-oxopregn-4-ene-20-carboxaldehyde (3-OPA). The deletion of this gene increases the production yield of the C-19 steroidal synthon 4-androstene-3,17-dione (AD) from natural sterols, avoiding the production of 4-HBC as by-product and the drawbacks in the AD purification. The molar yield of AD production using the MS6039-5941-6561 triple mutant strain was checked in flasks and bioreactor improving very significantly compared with the previously described MS6039-5941 strain.

## INTRODUCTION

One of the most attractive and efficient strategies for producing steroid precursors useful for the synthesis of steroid pharmaceutical drugs is to utilize mycobacterial strains to transform naturals sterols into useful intermediates or synthons as building blocks (Fernández-Cabezón *et al*., 2018; Donova, 2021; Nunes *et al*., 2022). Currently, three main C19 synthons are produced at industrial scale by different mycobacterial strains, i.e., 4-androstene-3,17-dione (AD), 1,4-androstadiene-3,17-dione (ADD) and 9α-hydroxy-4-androstenedione (9OH-AD) using phytosterols as natural precursors. C19 synthons are used in the chemical synthesis of sex hormones such as testosterone, estrone, estradiol and mineralocorticoids (Zhao *et al*., 2021; Feng *et al*., 2022). Apart from traditional mutation approaches, metabolic engineering techniques have been developed recently to manipulate the metabolism of sterols in these bacteria in order to accumulate the intermediate synthons at high yields in the culture medium (Fernández-Cabezón *et al*., 2018; Wang *et al*., 2022).

The aerobic sterol degradation pathway of mycobacterial strains has been studied extensively and most of the important key intermediates have been identified. It begins with the transport of sterols to the cellular interior mediated by the system, encoded by the *mce* operon (Pandey and Sassetti, 2008). Sterol degradation involves two main processes: the removal of the side chain and the cleavage of steroid rings. After the sterols have entered into the cell, two sequential reactions take place: the oxidation of cholesterol to 5-cholesten-3-one and its isomerization to 4-cholesten-3-one catalyzed by a 3β-hydroxysteroid dehydrogenase/isomerase (3β-HSD) that uses NAD or NADP as cofactors (Uhía *et al*., 2011). The aliphatic chain of sterols is degraded simultaneously, following a mechanism similar to β-oxidation of fatty acids, by three β-cyclic oxidations mediated by cytochromes P450 CYP125 and CYP142 (Capyk *et al*., 2009; Ouellet *et al*., 2010). Each cycle comprises enzymes such ligases, dehydrogenases, hydratases and one aldolase complex responsible of the AD formation and the release of two propionyl-CoA and one acetyl-CoA molecules (Fig. 1) (García *et al*., 2012). Considering this information, it has been demonstrated that the knockout of the *kstD, kshA* and *kshB* genes in different combinations allows the production of AD, ADD and 9OH-AD (Fernández-Cabezón *et al*., 2018; Wang *et al*., 2022).

**Figure 1.**
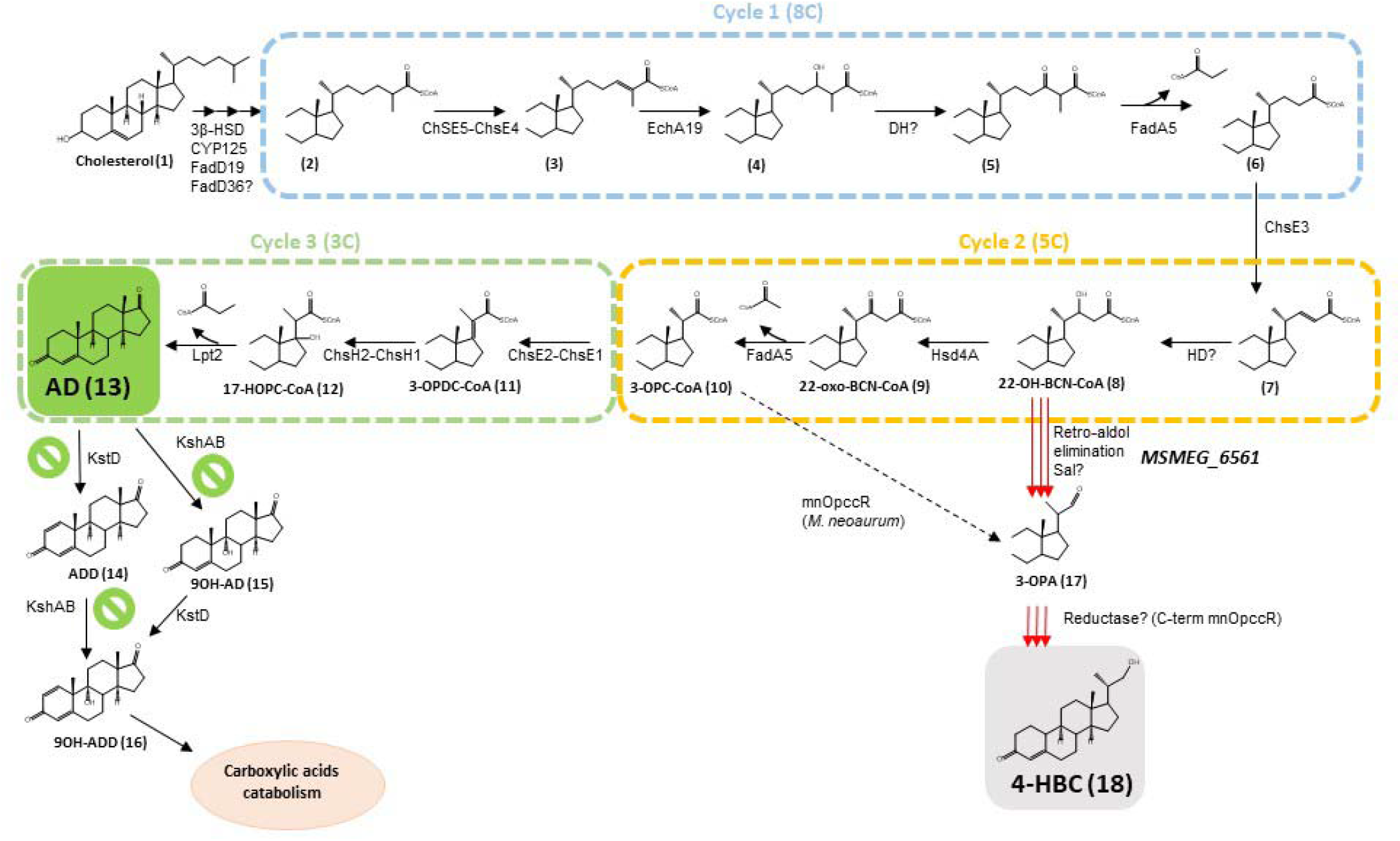
Cholesterol side-chain degradation pathway in Actinobacteria. Green symbols indicate the mutations performed to accumulate the C19 compounds. The three β-oxidation-like cycles are indicated with boxes. Enzymes with question marks are unknown and the biochemical steps have not been verified. Sal, steroid aldolase; DH, dehydrogenase; HD, hydratase. (1) Cholesterol, (2) (25S)-cholest-4-en-3-one-26-oyl-CoA, (3) Cholest-4,24-dien-3-one-26-oyl-CoA, (4) cholest-4-en-24-ol-3-one-26-oyl-CoA, (5) 3,24-dioxocholest-4-en-26-oyl-CoA, (6) 3-oxochol-4-en-24-oyl-CoA, (7) 3-oxochol-4,22-dien-24-oyl-CoA, (8) 22-hydroxyl-3-oxochol-4-en-24-oyl-CoA, (9) 3,22-dioxochol-4-en-24-oyl-CoA, (10) 3-oxo-23,24-bischornol-4-en-22-oyl-CoA, (11) 3-oxo-4,17-pregnadiene-20-carboxyl-CoA, (12) 3-oxo-23,24-bischornol-4-en-17-ol-22-oyl-CoA, (13) AD, (14) ADD, (15) 9OHAD, (16) 9OHADD, (17) 3-OPA, (18) 4-HBC.

Although well studied, there are still unknown aspects of *Mycolicibacterium* metabolism and in particular of *M. smegmatis* that need to be elucidated in order to achieve efficient industrial production of some intermediaries. For instance, the knockout of the *kstD (MSMEG_5941)* and *kshB (MSMEG_6039)* yielded an AD producer strain but it accumulates a certain amount of the by-product 4-HBC, which reduces the yield and hinders the purification of the final product (Galán *et al*., 2017).

Considering that *M. smegmatis* does not have an active mnOpccR reductase pathway that could explain the production of 4-HBC (Peng *et al*., 2021) (see below) (Fig.1), we propose that most probably this compound is produced in *M. smegmatis* by the aldolytic reaction to transform of 22-hydroxy-3-oxo-cholest-4-ene-24-carboxyl-CoA (22-OH-BCN-CoA) into (20S)-3-oxopregn-4-ene-20-carboxaldehyde (3-OPA) that is the proposed precursor of 4-HBC (Fig. 1). However, the enzymes involved in this transformation in *M. smegmatis* remained unknown. Retroaldol elimination of acetyl-CoA has been described in the degradation of the cholic acid side/chain in *Pseudomonas* sp. Chol1, renamed as *Stutzerimonas degradans* (Holert *et al*., 2013; Gomila *et al*., 2022). Both compounds, 22-OH-BCN-CoA, precursor of 4-HBC, and cholic acid are structurally similar and this reaction is catalyzed by Sal (steroid aldolase), a retro-aldolase (Holert *et al*., 2013).

In this work we have identified the *MSMEG_6561* gene encoding an aldolase enzyme responsible for the transformation of 22-OH-BCN-CoA into 3-OPA in *M. smegmatis*. We have demonstrated that a triple MS6039-5941-6561 mutant of *M. smegmatis* is unable to produce 4-HBC and therefore, it produces AD from natural sterols with a significant higher yield than the previous AD producer mutant.

## METHODS

### Chemicals

AD was purchased from TCI America. Chloroform and glycerol were purchased from Merck (Darmstardt, Germany). Cholesterol, cholestenone, N,O-bis(trimethylsilyl) trifluoroacetamide (BSTFA), gentamicin, pyridine, Tween 80 and tyloxapol were from Sigma (Steinheim, Germany). Oligonucleotides were from Sigma-Genosys. Phytosterols were provided by Gadea Biopharma (Spain) containing a mixture of different sterols (w/w percentage): brassicasterol (2.16%), stigmasterol (8.7%), campesterol (36.8%) and β-sitosterol (54.4%).

### Bacterial strains and culture conditions

The strains as well as the plasmids and primers used in this work are listed in Table 1. Preinocula of *M. smegmatis* mc^2^ 155 and its derivative strains were grown in LB medium with 0.05% Tween 80 (Sigma) at 37°C with orbital shaking at 200 rpm. LB agar plates and 7H10 agar (Difco) with sucrose 10% (w/v) were used for solid media. *E. coli* strains were grown in LB medium at 37°C with orbital shaking at 200 rpm and in LB agar plates for solid media.

**Table 1.**
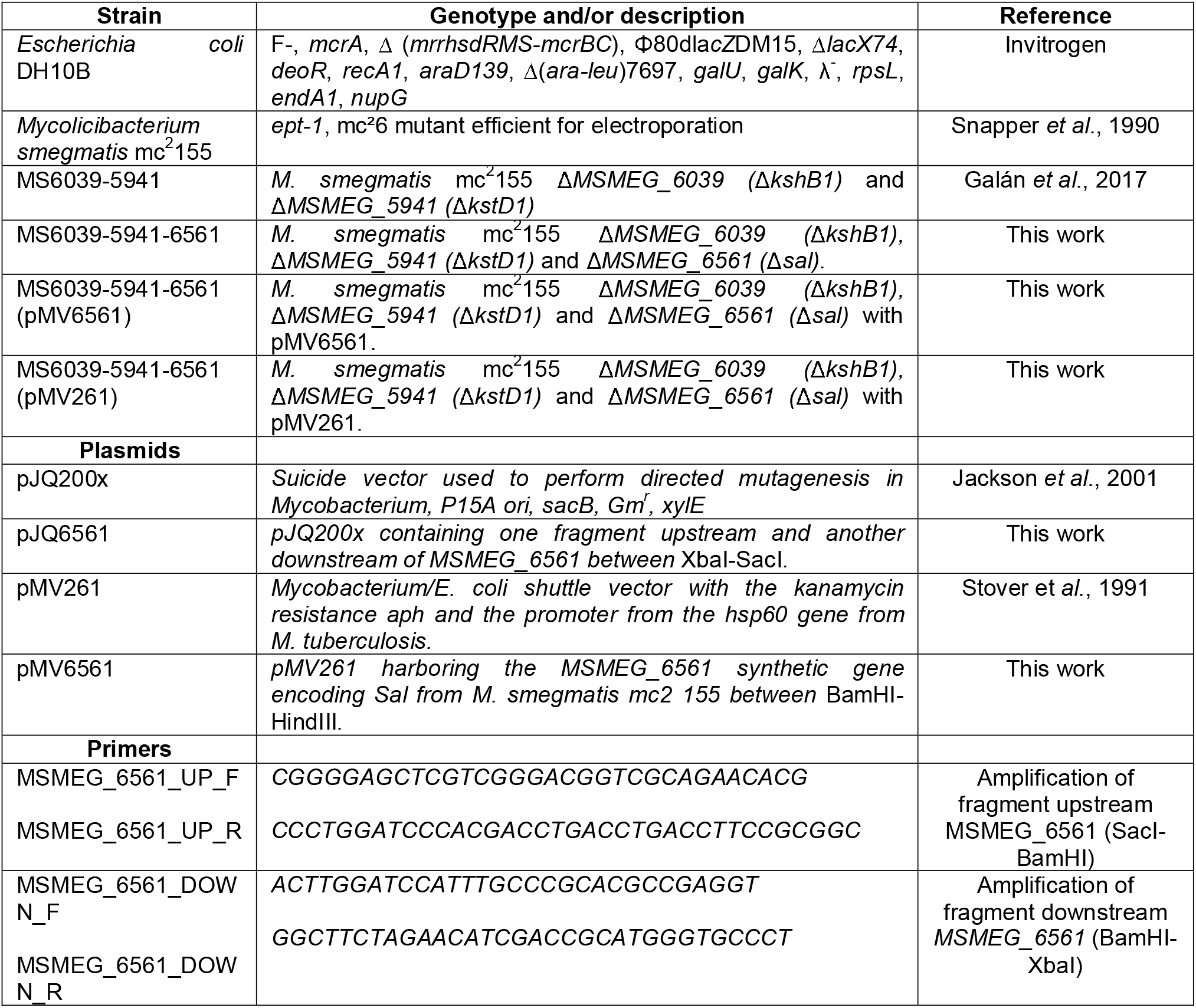
Strains, plasmids and primers

For the biotransformation assays in 100 ml flasks, Middlebrook 7H9 broth medium (Difco) supplemented with 9 mM glycerol (Sigma) as carbon and energy source and 0.4 g/l cholesterol (Sigma) as raw material were used. Due to the high hydrophobicity of cholesterol, the 5 mM stock was prepared in tyloxapol 10% (v/v) (Sigma), sonicated for 30 min and autoclaved twice until its total dissolution. Tyloxapol final concentration in the biotransformation medium was 2%.

The preinoculum was first grown over-weekend on LB with Tween 80 as described before. Then, it was washed twice with saline solution 0.85% with 0.05% Tween 80 and the biotransformation assays were inoculated at *OD*_*600*_=0.05 in 25 ml of biotransformation media. Strains were grown at 200 rpm and 37°C for 96 h and 500 μl samples of the culture were taken every 24 h and kept at -20°C until their analysis by GC/MS.

Bioreactor experiments were performed using the culture media previously described (Kutney *et al*., 2003) containing: corn steep liquor (20 g/l), sodium nitrate (5.4 g/l), ammonium chloride (0.28 g/l) and monobasic sodium phosphate (0.28 g/l). 10 g of phytosterols were dissolved in 160 ml of sunflower oil and mixed with the aqueous phase (740 ml). The inoculum volume was 10%. This experiment was carried out in a commercial stirred tank bioreactor (STBR) of 2 l total volume (Sartorius AG, Germany) and a working volume of 1 l was used. The process was carried out at 37 °C, pH was adjusted at the start of the reaction to 7.4 and was not controlled during the process, 600 rpm of stirrer speed and an air flow rate of 1 vvm were used. Dissolved oxygen was maintained above 20% throughout the process. When needed, final concentrations of 5 μg/ml of gentamicin and 20 μg/ml of kanamycin were added to the growth media for *M. smegmatis* strains, while 10 μg/ml of gentamicin and 50 μg/μl of kanamycin were used for the *E. coli* strains.

### Gene deletion

The knock-out strain of *M. smegmatis* named MS6039-5941-6561 was constructed by homologous recombination using the pJQ200x plasmid, a derivative of the suicide pJQ200 vector that does not replicate in *Mycobacterium* (Jackson *et al*., 2001). The strategy consist in generating two fragments of ~600 bp, the first one containing the upstream region and few nucleotides of the 5′-end of the gene and the second one containing the downstream region and few nucleotides of its 3′-end of the gene, that are amplified by PCR using the oligonucleotides described in Table 1 and *M. smegmatis* genomic DNA as template. *M. smegmatis* genomic DNA extraction was performed as described (Uhía *et al*., 2011). The two fragments generated were digested with the corresponding enzymes and cloned into the plasmid pJQ200x using *E. coli* DH10B competent cells as described (Uhía *et al*., 2011). Plasmid DNA from *E. coli* DH10B recombinant strain was extracted using the High Pure Plasmid Purification Kit (Roche, Basel, Switzerland), according to the manufacturer’s instructions. This procedure was performed for gene *MSMEG_6561* generating the pJQ6561plasmid. Plasmid pJQ6561 was electroporated into competent *M. smegmatis* MS6039-5941 to obtain the strain MS6039-5941-6561. Single cross-overs were selected on 7H10 agar plates containing gentamicin and the presence of the *xylE* gene encoded in pJQ200x was confirmed by spreading catechol over the single colonies of electroporated *M. smegmatis*. The appearance of a yellow coloration indicates the presence of the *xylE* gene. Colonies were also contra-selected in 10% sucrose. A single colony was grown in 10 ml of LB medium with 5 μg/ml gentamicin up to an optical density of 0.8–0.9 and 20 μl of a 1:100 dilution was plated onto 7H10 agar plates with 10% sucrose to select for double cross-overs. Potential double cross-overs (sucrose-resistant colonies) were screened for gentamicin sensitivity and the absence of the *xylE* gene. The mutant strain MS6039-5941-6561 was analyzed by PCR and DNA sequencing to confirm the deletions of *MSMEG_6561* gene.

### Construction of pMV6561 plasmid

*MSMEG_6561* gene from *M. smegmatis* mc^2^ 155 genomic DNA was chemically synthesized by GeneScript (The Netherlands). The synthetic gene was digested with BamHI and HindIII and cloned into pMV261, a shuttle plasmid that replicates in *E. coli* and *M. smegmatis* generating the plasmid pMV6561 that was transformed into *E. coli* DH10B to generate the recombinant strain *E. coli* DH10B (pMV6561). Once the sequence of the pMV6561 was checked it was used to transform electrocompetent cells of *M. smegmatis* MS6039-5941-6561 generating the *M. smegmatis* MS6039-5941-6561 (pMV6561) recombinant strain.

### GC/MS analyses

To perform GC/MS analysis, culture aliquots (0.2 ml) were extracted twice at various extents of incubation with an equal volume of chloroform. Previously to its extraction, 10 μl of a solution of 10 mM cholesterol (when phytosterols were used as substrate) or 10 mM cholestenone (when cholesterol was used as substrate) dissolved in chloroform were added to the aliquots as internal standards. The chloroform fraction was concentrated by evaporation and the trimethylsilyl ether derivatives were formed by reaction with 50 μl of TMT-BSTFA and 50 μl of pyridine and heating at 80°C for 20 min. Calibration standards were derivatized in the same way. The GC/MS analysis was carried out using an Agilent 7890A gas chromatograph coupled to an Agilent 5975C mass detector (Agilent Technologies, Palo Alto, CA, USA). Mass spectra were recorded in electron impact (EI) mode at 70 eV within the m/z range 50–550. The chromatograph was equipped with a 30 m x 0.25 mm i.d. capillary column (0.25 μm film thickness) HP-5MS (5% diphenyl 95% dimethylpolysiloxane from Agilent Technologies). Working conditions in the sample were as follows: split ratio (20:1), injector temperature, 320°C; column temperature 240°C for 3 min, then heated to 320°C at 5°C min−1. For quantification of the peak area, the quantitative masses were 329 + 458 m/z for cholesterol; 343 + 384 m/z for cholestenone; 382 + 472 m/z for campesterol; 394 + 484 m/z for stigmasterol; 357 + 486 m/z for β-sitosterol; 244 + 286 m/z for AD and 73 + 270 m/z for 4-HBC in selected ions of monitoring. EI mass spectra and retention data were used to assess the identity of compounds by comparing them with those of standards in the NIST Mass Spectral Database and commercial standards (NIST 2011).

### Calculations

The molar yield (MY) of the products were determined according to the following equation [Eq. 1]:

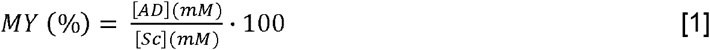

In the equation, the term “AD” refers to 4-androstene-3,17-dione and “Sc” refers to consumed sterols.

The productivity (P) of 4-androstene-3,17-dione (AD) was determined according the following equation (Eq. 2):

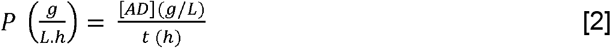

In the equation, the term “t” refers to time (h) and “Sc” refers to consumed sterols.

## RESULTS AND DISCUSSION

### Identification of the aldolase-encoding gene in the genome of *M. smegmatis*

Mutants constructed to obtain C19 synthons accumulate C22 intermediate by-products, reducing yield and making purification of the final product difficult (Galán *et al*., 2017). In the same way, mutants with the blocked side chain degradation pathway constructed to produce C22 steroids show accumulation of C19 residues (Xu *et al*., 2016; Galán *et al*., 2017). Thus, looking to the side-chain sterol degradation pathway we can found a conventional route by which AD and other C19 steroids are generated as degradation intermediates, and an alternative route that gives rise to 4-HBC and other C22 steroids as final degradation products. The central node of both pathways is 22-OH-BCN-CoA, which leads to AD or to 4-HBC through two possible pathways, one proposed in *Mycolicibacterium neoaurum* (Peng *et al*., 2021) and an unknown aldolytic reaction (Fig.1).

The pathway described in *M. neoaurum* starts from 3-oxo-4-pregnen-20-carboxyl-CoA (22-OPC-CoA) that is generated from 22-OH-BCN-CoA by the consecutive action of two enzymes, *i. e*., Hsd4A (a dual-function enzyme, with both 17β-hydroxysteroid dehydrogenase and β-hydroxyacyl-CoA dehydrogenase activities) (Xu *et al*., 2016) and FadA5 thiolase (Nesbitt *et al*., 2010) (Fig. 1). Next the transformation of 22-OPC-CoA into 4-HBC is made by means of a dual function reductase named mnOpccR. mnOpccR acts catalyzing both a 4e-reduction reaction and a 2e-reduction reaction for the formation of 4-HBC. For the 4e-reduction, a first 2e-reduction takes place at the C-terminus of the enzyme, to convert the 3-OPC-CoA to 3-oxo-4-pregnen-20-carboxyl aldehyde (3-OPA) followed by a second 2e-reduction, at the N-terminus, of the aldehyde group to alcohol to form 4-HBC. In the 2e-reduction reaction, the only substrate is 3-OPA, which is reduced at the N-terminus of mnOpccR. It has been shown that inactivation or overexpression of mnOpccR in *M. neoaurum* can achieve exclusive production of AD or 4-HBC (Peng *et al*., 2021).

In *M. smegmatis*, a gene homologous to *mnopccR* was found with 78% identity, i.e., *MSMEG_1623* (accession number: AIU06854). However, Peng *et al*. (2021) have demonstrated that the C-terminal end of the MSMEG_1623 protein is not active. Therefore, the only way to obtain 4-HBC in *M. smegmatis* would be through the degradation of the 22-OH-BCN-CoA through an alternative route. This conversion could occur via a retroaldol elimination, or aldolytic cleavage, that would produce 3-OPA (Fig. 1), followed by a reduction by the N-terminal domain of msOpccR reductase to generate 4-HBC (Peng *et al*., 2021) or by another reductase activity.

The retroaldol elimination of acetyl-CoA has been described in the degradation of the cholic acid side chain in *S. degradans* (Holert *et al*., 2013). Both compounds, 22-OH-BCN-CoA, precursor of 4-HBC, and cholic acid are structurally similar. This reaction is catalyzed by the enzyme called Sal (steroid aldolase), encoded by the *sal* gene (Holert *et al*., 2013). Chemically, the aldolytic reaction of 22-OH-BCN-CoA (24C) to 3-OPA (22C aldehyde) could be performed by a *sal*-encoded aldol-lyase similar to that of the degradation of cholate side chain in *S. degradans*.

To identify the gene responsible for encoding this activity in *M. smegmatis*, we searched using Blastp those sequences similar to the Sal enzyme within its genome. Thus, the *MSMEG_6561* gene was identified, whose amino acid sequence shares a 57% similarity with the Sal enzyme (OOE10244.1) of *S. degradans*. Other *Mycolicibacterium* such as *M. neoaurum* or *M. fortuitum* that has been used for the production of steroid synthons also encode a homologous Sal enzyme (Table 2). Moreover, *Mycobacterium tuberculosis* has also a homologous protein (Table 2).

**Table 2.**
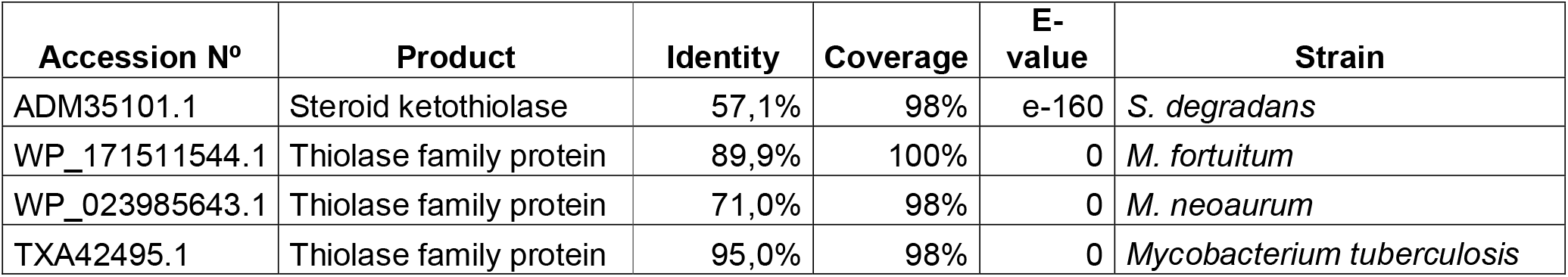
Proteins homologous to MSMEG_6561.

| Accession Nº | Product | Identity | Coverage | E-value | Strain |
| --- | --- | --- | --- | --- | --- |
| ADM35101.1 | Steroid ketothiolase | 57,1% | 98% | e-160 | *S. degradans* |
| WP_171511544.1 | Thiolase family protein | 89,9% | 100% | 0 | *M. fortuitum* |
| WP_023985643.1 | Thiolase family protein | 71,0% | 98% | 0 | *M. neoaurum* |
| TXA42495.1 | Thiolase family protein | 95,0% | 98% | 0 | *Mycobacterium tuberculosis* |

Remarkable, the *MSMEG_6561* gene is expressed constitutively in *M. smegmatis* (Uhía *et al*., *2012)*. The transcriptomic analysis carried out in the presence or absence of cholesterol indicates that *MSMEG_6561* is not induced in the presence of cholesterol suggesting that it could play other role in the metabolism of some related compounds.

*MSMEG_6561* gene is part of a putative operon containing *MSMEG_6560*, encoding a putative enoyl-CoA-hydratase, *MSMEG_6562*, encoding a putative acyl-CoA dehydrogenase, and *MSMEG_6563* coding a putative oxidoreductase having Rossmann-fold NAD(P)-binding domain-containing protein according to the Conserved Domain DB at NCBI. The resemblance of this operon to that of the *S. degradans* strain, where the aldolase Sal and hydratase Shy coding genes are located contiguous in the genome (Holert *et al*., 2013), suggests that it could very likely to be involved in the formation of 3-OPA and subsequent reduction to 4-HBC in the MS6039-5941 mutant strain. Elucidation of the specific role of this operon in the metabolism of *M. smegmatis* requires further studies but most probably could be involved in the degradation of some related compounds such bile acids and/or fatty acids.

### Construction of the triple MS6039-5941-6561 mutant

To study the possible aldolase activity of the *MSMEG_6561* gene, we have mutated the strain *M. smegmatis* MS6039-5941 (Δ*kshB1*, Δ*kstD1*) which is the AD producer that accumulates 4-HBC as a by-product (Galán *et al*., 2017) by double homologous recombination yielding the triple mutant *M. smegmatis* MS6039-5941-6561 (Δ*kshB1*, Δ*kstD1*, Δ*sal*). The mutation of the *MSMEG_6561* gene was confirmed by PCR and DNA sequencing by Sanger.

The MS6039-5941-6561 mutant is able to grow in minimal medium using glycerol or glucose as carbon and energy sources as efficiently as the MS6039-5941 mutant, indicating that the mutation of *MSMEG_6561* gene does not affect the growth in these substrates (data not shown). In addition, the morphology of the mutant cells and the plate colonies does not appear to be affected.

### Production of AD in the *M. smegmatis* MS6039-5941-6561 strain

The mutant strains MS6039-5941 and MS6039-5941-6561 were grown in minimal medium with glycerol as energy and carbon sources and in the presence of 0.4 g/l cholesterol to address the production of steroid intermediates in flasks. Figure S1 shows that the triple mutation has not impaired strain growth in the biotransformation medium compared with the MS6039-5941 strain. Steroids were extracted from aliquots of the culture medium every 24 h for 96 h and were separated and identified by GC/MS, as detailed in Materials and Methods. As previously mentioned, the degradation of the sterol side chain can follow two possible routes in *M. smegmatis*: the conventional one by which AD and other C-19 steroids are generated as degradation intermediates and an alternative route that gives rise to the 4-HBC and other C-22 steroids as degradation end products (Fig. 1). Figure 2 show the chromatograms and mass spectra obtained from the samples of MS6039-5941 and MS6039-5941-6561 strain, respectively, taken at 96 h. While in the case of MS6039-5941 strain, three major peaks are observed: (1 and 2) corresponding to the accumulated products: AD and 4-HBC, respectively, and (4) corresponding to 4-cholesten-3-one (internal standard, hereinafter cholestenone). There is not 4-HBC detected when the triple mutant strain was used. Other minor peaks that have been identified as non-steroidal trimethylsilyl residues of the reagents used in the derivatization of the samples are observed. MS6039-5941-6561 strain reached up to 240 mg/l of AD after 96 h, while MS6039-5941 only produced up to 150 mg/l (Fig. 3). Molar yields of the biotransformation process after 96 h reached 59% and 81% for the MS6039-5941 and MS6039-5941-6561 strains, respectively (Fig. 3).

**Figure 2.**
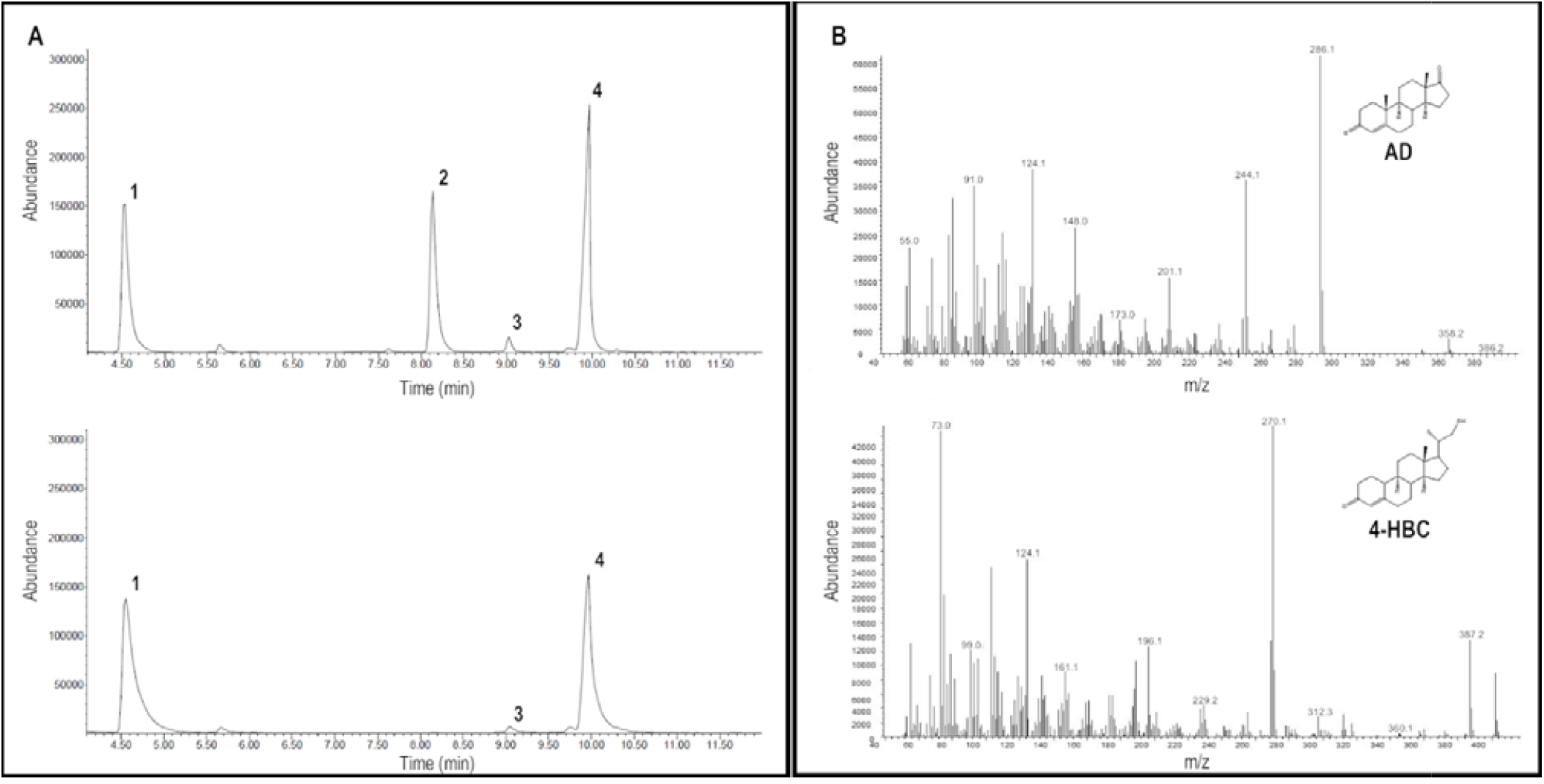
A) Analysis by GC/MS of the products accumulated in the culture of MS6039-5941 (up) and MS6039-5941-6561 (down) strains after 96 h in a medium containing 9 mM glycerol and cholesterol at 0.4 g/l. (1) AD, (2) 4-HBC, (3) cholesterol and (4) cholestenone (internal standard). B) Mass spectra and fragmentation pattern of AD (up) and 4-HBC (down).

**Figure 3.**
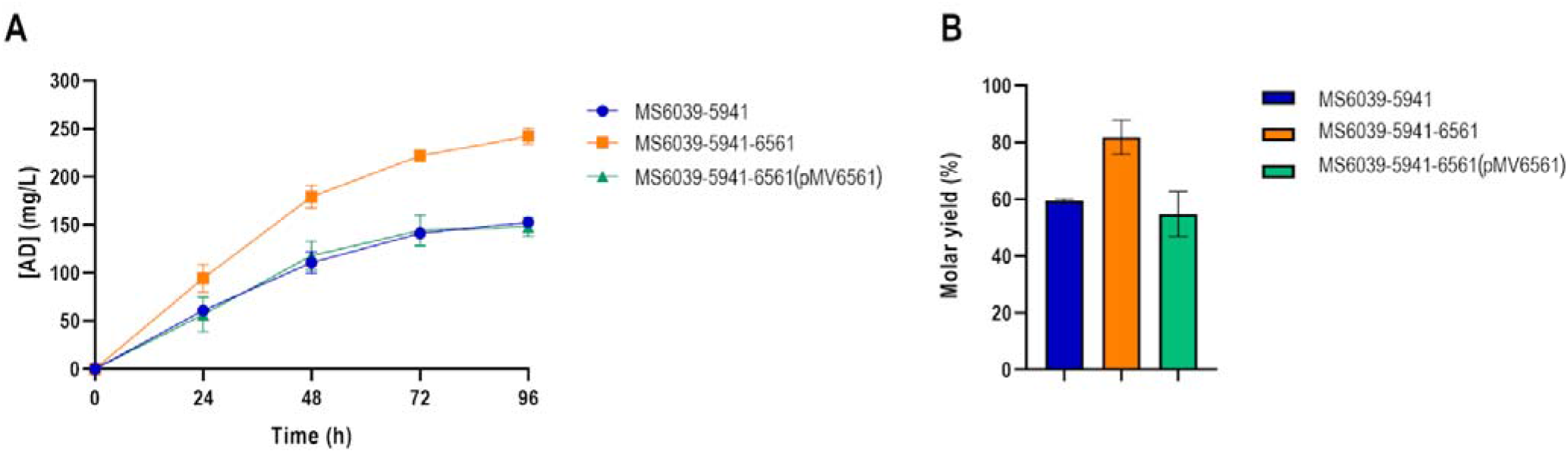
A) AD production of the MS6039-5941, MS6039-5941-6561 and MS6039-5941-6561 (pMV6561) strains cultured on shake flasks in a medium containing 9 mM glycerol and cholesterol at 0.4 g/l. B) Molar yield of AD production of MS6039-5941, MS6039-5941-6561 and MS6039-5941-6561 (pMV6561) strains.

To further confirm that *MSMEG_6561* is the coding gene responsible of the accumulation of 4-HBC during the cholesterol biotransformation, the MS6039-5941-6561 strain was transformed with plasmid pMV6561 carrying the *MSMEG_6561* gene under the control of P_*hsp60*_ promoter yielding the MS6039-5941-6561 (pMV6561) strain (Table 1). The resulting strain was checked in a biotransformation assay in the presence of 0.4 g/l of cholesterol. First, the ability to grow in the biotransformation medium at the same rate than the parental strain was confirmed (Fig. S1). The GC-MS analysis of the supernatant show that the complementation with *MSMEG_6561* gene allows the formation of the by-product 4-HBC (Fig. S2). MS6039-5941-6561 (pMV6561) strain showed the same levels of AD production as MS6039-5941 (around 150 mg/l) and the molar yield after 96 h reached 60% (Fig. 3)

To determine the behavior of the new strain using phytosterols as a substrate, a biotransformation assay was carried out in a STBR. The MS6039-5941-6561 mutant was grown for 96 h and samples were taken every 24 h. Figure 4A shows the phytosterol consumption and the AD production over time by this strain. Remarkably, the new strain was able to produce up to 7 g/l of AD. In these conditions, a productivity of 0.17 gl^-1^ h^-1^ of AD is reached at 24 h and a productivity of 0.12 gl^-1^ h^-1^ at 48 h, indicating that the maximum production of AD occurs in the first 24 h. Moreover, molar yield of this biotransformation after 72 h reached 98% (Fig. 4B). After 96 h we have not detected the presence of 4-HBC and only traces of non-consumed stigmasterol can be detected by GC-MS (Fig. S3). The aliphatic side-chain of stigmasterol contains a double bond that might explain why this compound is metabolized less efficiently than cholesterol, campesterol or β-sitosterol. In this sense, β-oxidation of side-chain of cholesterol or phytosterols is not identical (Chen *et al*., 1985; Fujimoto *et al*., 1982b; Wilbrink *et al*., 2012). Some biochemical steps are described to be specific for phytosterols side-chain degradation that in turn are not essential for cholesterol, like the CoA activation by FadD19 and the retro-aldol elimination of propionyl CoA by Ltp3 and/or Ltp4 aldol lyases occurring in the first β-oxidation cycle (Wilbrink *et al*., 2012; Wronska *et al*., 2016). However, further CoA activation steps, are followed by two rounds of β-oxidation and formation of a C19 steroid via aldolytic cleavage are identical to cholesterol side-chain degradation. Thus, it is becoming clear that phytosterols are substrates that require a more specific arsenal of enzymes than cholesterol for the degradation of the branched side-chain. Therefore, deepening the knowledge of the enzymes involved in this process will allow us to build more efficient production strains.

**Figure 4.**
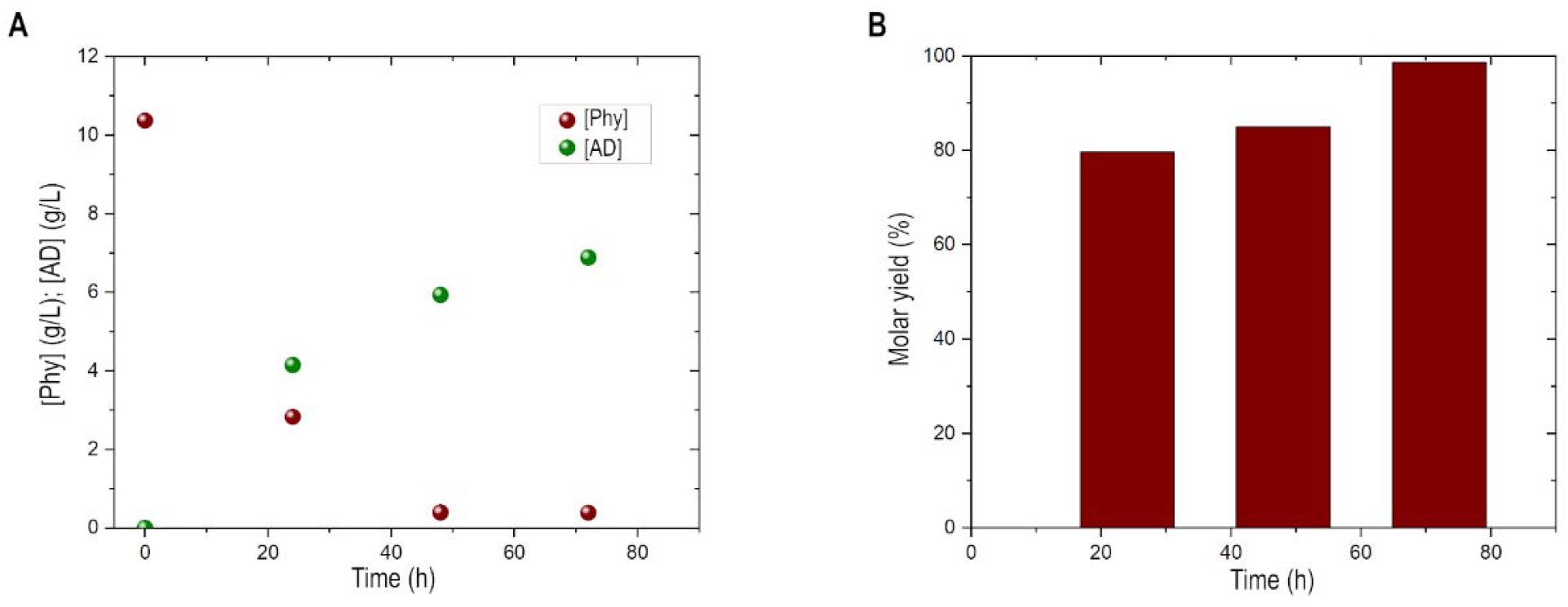
A) AD production and phytosterols consumption over time of the MS6039-5941-6561 strain cultured in a bioreactor containing phytosterols at 10 g/l. B) Molar yield of AD production of the MS6039-5941-6561 strain.

## CONCLUSION

The results presented in this work demonstrate that the *MSMEG_6561* gene encoding a putative aldolase is involved in the production of 4-HBC in the metabolism of sterols in *M. smegmatis*. This gene is homologous to the *salI* gene of some bacteria able to degrade bile salts. In *M. smegmatis* the gene appears to form part of an operon involved in a β-oxidation process that is not necessary for cholesterol degradation.

Since 4-HBC cannot be used by *M. smegmatis* as a carbon and energy source, the presence of this end-product can only be explained because the blockage of the pathway generates an accumulation of CoA-intermediaries that are not efficiently processed by the sterol site-chain β-oxidation enzymes. When *M. smegmatis* is cultured on high concentrations of sterols, the accumulation of CoA-derivatives can be toxic for the cells since it reduces the availability of CoA for other processes hindering the bacterial growth. In these cases, thiolases or aldolases having a broad range substrate specificity can help to release CoA from the overproduced CoA-derivatives. The release of CoA is concomitant to the production of acetyl-CoA that can be readily metabolized.

It is worth to notice that, in contrast to the metabolism of bile-salts, where the resulting aldehyde is oxidized to a carboxyl group by the aldehyde dehydrogenase Sad (Holert *et al*., 2013), in *M. smegmatis* the aldehyde is reduced to an hydroxylic group.

Although we cannot completely discard that other genes could play a similar role to that of *MSMEG_6561*, at least this gene appears to be the main responsible for the production of 4-HBC in *M. smegmatis* under the tested conditions and in the absence of an active msOpccR reductase (Fig. 1). Interestingly, *sal* gene does not appear to play the same critical role in *M. neoaurum* since the deletion of *msopccR* was enough to prevent the production of 4-HBC (Peng *et al*., 2021), therefore the role of the homologous *sal* genes in other *Mycolicibacterium* should be further investigated.

## AUTHOR CONTRIBUTION

Gabriel Hernández-Fernández: Formal analysis (supporting); investigation (equal); methodology (equal); writing – review and editing (supporting). Miguel G. Acedos: Formal analysis (supporting); investigation (equal); methodology (equal); writing – review and editing (supporting). José L. García Conceptualization (lead); formal analysis (lead); funding acquisition (lead); writing – review and editing (lead). Beatriz Galán: Conceptualization (lead); formal analysis (lead); funding acquisition (lead); writing – review and editing (lead).

## ACKNOWLEDGMENTS

This work was supported by grants from Spanish Ministry of Science and Innovation PID2021-125370OB-I00, PID2019-110612RB-I00 from the Ministry of Economy and Competitiveness of Spain and grant from the Community of Madrid and the Structural Funds from European Union (Ref: S2018/BAA-4532 (ALGATEC-CM)). The grant for one of the authors (GHF) with reference FPU21/05101 is gratefully recognized. The authors would like to thank Ana Valencia for the technical assistance.

## Supplementary Material

**Figure S1.**
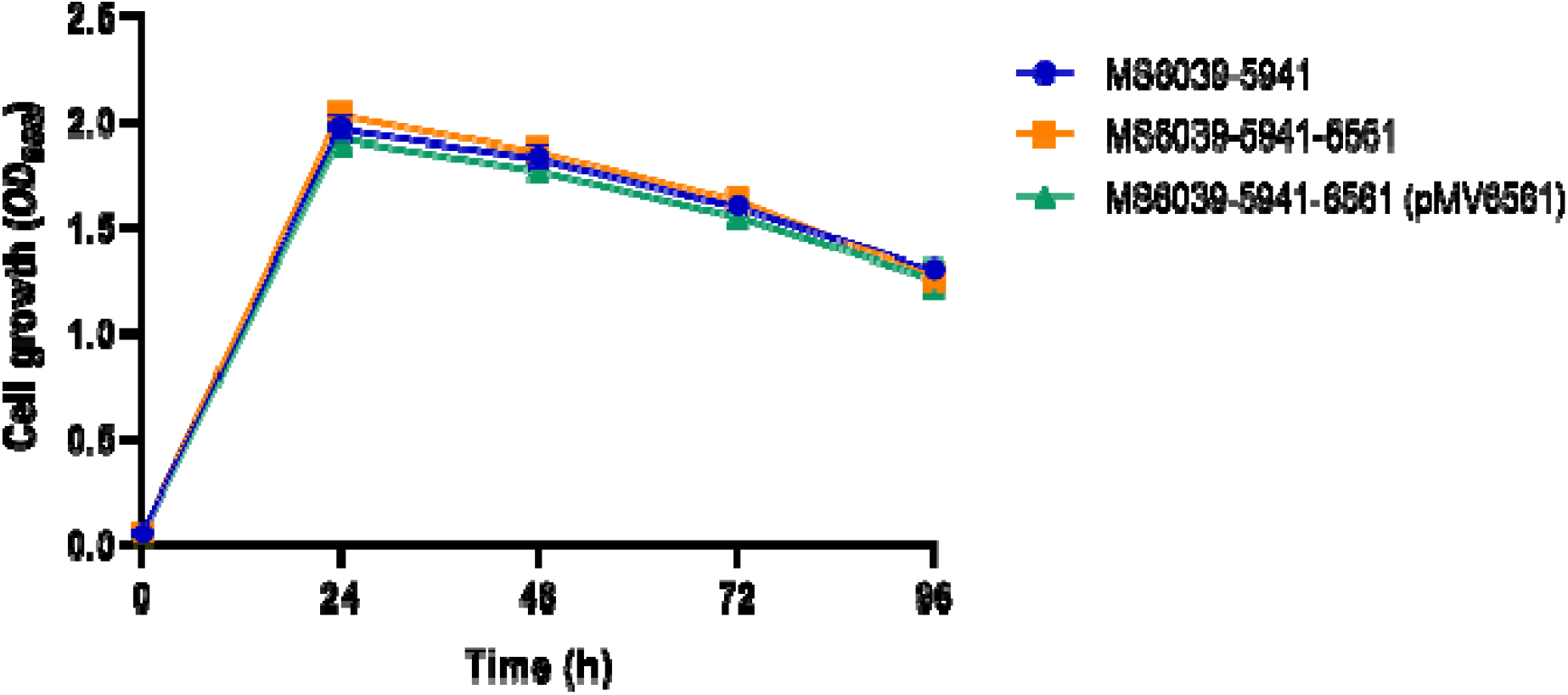
Growth curve of MS6039-5941, MS6039-5941-6561 and MS6039-5941-6561 (pMV6561) in shaken flasks on the biotransformation medium containing 9 mM glycerol and cholesterol at 0.4 g/l.

**Figure S2.**
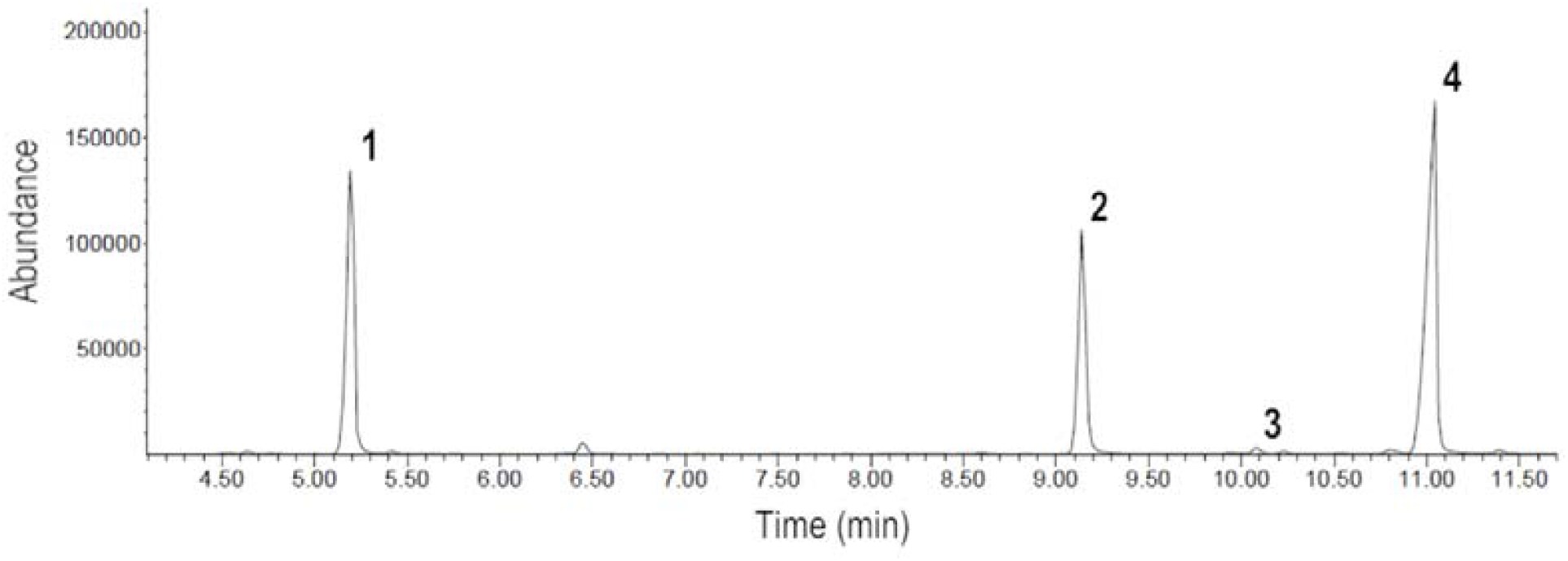
Analysis by GC/MS of the steroids accumulated in the culture of MS6039-5941-6561 (pMV6561) strain after 96 h in shaken flasks on a medium containing 9 mM glycerol and cholesterol at 0.4 g/l. (1) AD, (2) 4-HBC, (3) cholesterol and (4) cholestenone (internal standard).

**Figure S3.**
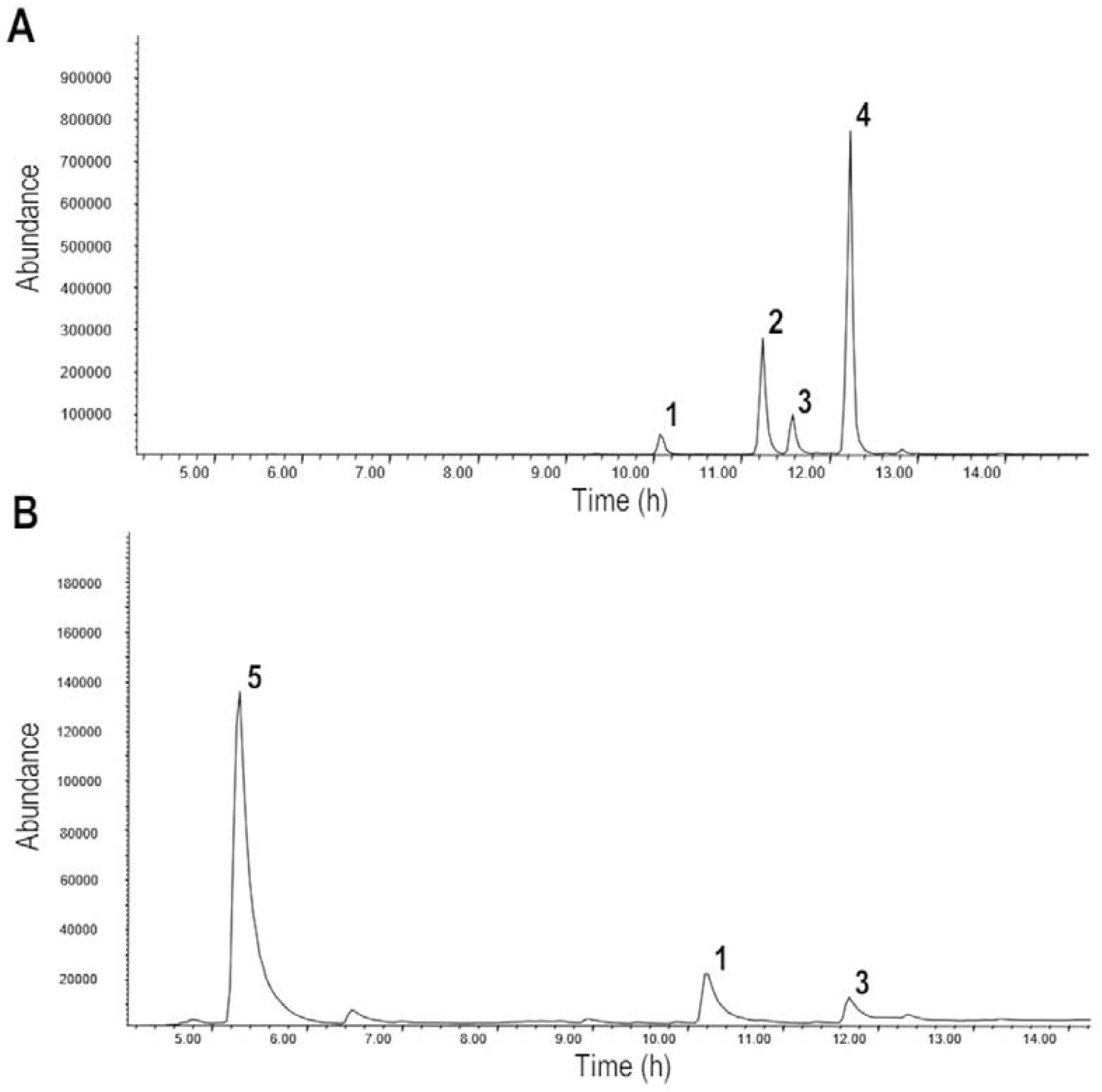
Analysis by GC/MS of the steroids accumulated in the culture of MS6039-5941-6561 strain after 96 h in a bioreactor containing phytosterols at 10 g/l. A) Chromatogram at 0 h. B) Chromatogram at 96 h. (1) cholesterol (internal standard), (2) campesterol, (3) stigmasterol, (4) β-sitosterol and (5) AD.

